# Dynamic hippocampal-cortical interactions during event boundaries support retention of complex narrative events

**DOI:** 10.1101/2022.10.23.513391

**Authors:** Alexander J. Barnett, Mitchell Nguyen, James Spargo, Reesha Yadav, Brendan I. Cohn-Sheehy, Charan Ranganath

## Abstract

According to most memory theories, encoding involves continuous communication between the hippocampus and neocortex leaving the temporal dynamics of hippocampal-neocortical interactions often overlooked. Recent work has shown that we perceive complex events in our lives as dynamic, with relatively distinct starting and stopping points known as event boundaries. Event boundaries may be important for memory, as they are associated with increased activity in the hippocampus, and extended neocortical regions (the posterior cingulate cortex, lateral parietal cortex, and parahippocampal cortex). Our objective was to determine how functional connectivity between the hippocampus and neocortical regions during the encoding of naturalistic events (movies) related to subsequent retrieval and retention of those events. Participants encoded two 16-minute cartoon movies during fMRI scanning. After encoding, participants freely recalled one of the movies immediately, and the other after a 2-day delay. We quantified hippocampal-neocortical functional connectivity (FC) at time windows around each event onset, middle, and offset, and compared these FC measures with subsequent recall. These analyses revealed that higher FC between the hippocampus and the posterior medial network (PMN) at an event’s offset related to whether that event was subsequently recalled. In contrast, mid-event connectivity between the hippocampus and PMN was associated with poorer memory. Furthermore, hippocampal-PMN offset connectivity predicted not only whether events were retained in memory, but also the degree to which these events could be recalled in detail after a 2-day delay. These data demonstrate that the relationship between memory encoding and hippocampal-neocortical interaction is more dynamic than suggested by most memory theories, and they converge with recent modeling work suggesting that event offset is an optimal time for encoding.

---

Since the pioneering work of Ebbinghaus (Ebbinghaus, 1885), researchers have studied memory by investigating recognition or recall of lists of verbal stimuli or pictures. Neuroscience research in this tradition has shown that memory for this kind of arbitrary episodic information is supported by the hippocampus, which is thought to index neocortical representations during an event (Frankland and Bontempi, 2005; McClelland et al., 1995; Nadel and Moscovitch, 1997; Squire and Zola-Morgan, 1991; Teyler and DiScenna, 1986). Real-world events, however, are not entirely arbitrary. Considerable evidence suggests that we can use our prior knowledge of event structure to generate mental models (called event models) that enable inferences about current experiences and prediction of upcoming information (Radvansky and Zacks, 2014; Rumelhart and Ortony, 1977; Thorndyke, 1977). It is our knowledge of this predictive structure that led Bartlett to note that memories for events in our lives are not an exact replay of past events, but rather are an “imaginative reconstruction” of the past based on our prior knowledge of the world (Bartlett, 1932). Different theories have been proposed to explain memory for arbitrary event information (Cohen et al., 1997; Howard et al., 2005), compared to constructive elements of memory and event cognition (Radvansky and Zacks, 2014; Rumelhart and Ortony, 1977).

Recent work investigating encoding and retrieval of structured events in films and stories (Bird, 2020; Finn, 2021; Hasson et al., 2015; Lee et al., 2020) has indicated, surprisingly, that the default mode network (DMN) carries event-specific information in multivariate fMRI activation patterns during event encoding and retrieval (Baldassano et al., 2017; Chen et al., 2017; Oedekoven et al., 2017; Reagh and Ranganath, 2021). These regions tend to have stable activation patterns within events and shift their patterns at points that coincide with subject-identified event boundaries (Baldassano et al., 2017; Ben-Yakov and Henson, 2018; Geerligs et al., 2021). Hippocampal activity, on the other hand, increases during transitions between events, also known as event boundaries (Reagh et al., 2020; Zheng et al., 2022), which in turn, coincide with pattern shifts in the DMN (Baldassano et al., 2017). This hippocampal activity at the end of an event is reliably correlated with successful encoding of event details (Ben-Yakov and Dudai, 2011; Ben-Yakov et al., 2013).

Recent cognitive modeling has suggested that event information, represented in the neocortex, is encoded into the hippocampus and this may preferentially occur at event offsets (Franklin et al., 2020; Lu et al., 2022). The prediction that the hippocampus disproportionately indexes neocortical activity at event boundaries is a radical departure from traditional models of memory, but it does align with the Complementary Learning Systems framework which suggests that the hippocampus may be specialized for rapid encoding of arbitrary associations (i.e., information at event boundaries) whereas neocortical networks may be optimized for learning about regularities across events (i.e., event schemas)(McClelland et al., 1995; O’Reilly et al., 2022).

The neocortex, however, is not a homogeneous collection of regions and specific networks of neocortical regions are thought to play a role in representing event models and connect with the hippocampus. Analyses of cortical networks with fMRI have indicated that the hippocampus shows high functional connectivity with a “Medial Temporal Network” (MTN) and 3 subnetworks of the DMN (Andrews-Hanna et al., 2010; Barnett et al., 2021; Cooper et al., 2021; Gordon et al., 2020)—the “Posterior Medial Network” (PMN), “Anterior Temporal Network (ATN)”, and the “Medial Prefrontal Network” (MPN). These networks can be differentiated based on multivariate representations during memory-guided decision making (Barnett et al., 2021), task activation (DiNicola et al., 2020), and functional connectivity patterns (Barnett et al., 2021; Braga et al., 2019; Cooper et al., 2021; Gordon et al., 2020). While these networks show reliable connectivity with the hippocampus, it is unclear how their interactions with the hippocampus support encoding and whether this support is amplified at event boundaries.

Here, using fMRI, we tested the idea that the encoding of structured events is supported by cortico-hippocampal interactions at event boundaries. Participants were scanned while viewing custom-made animated films and subsequently recalled events from these films either in the same scanning session, or after a 2-day delay (Figure 1A). We tested the prediction that functional connectivity between the hippocampus and neocortical will be associated with subsequent retrieval success and this association would be specifically pronounced at the offset of an event. Based on evidence that recall quality of naturalistic events changes over time (Fisher and Radvansky, 2018; Sekeres et al., 2016, 2018), we also examined whether functional interactions at encoding were particularly important for detailed memory after a longer (2-day) versus shorter (5 minute) delay.

**Figure 1.**
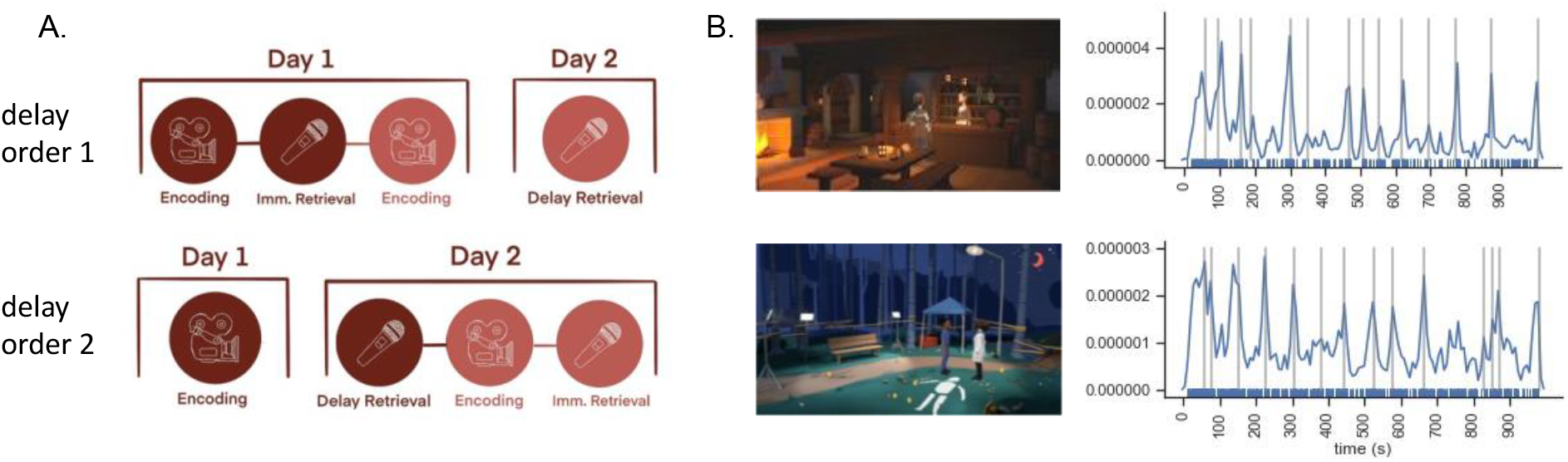
A. overview of procedure. B. screen shot from each movie with accompanying kernel density estimates of event boundaries from 62 online participants. Gray Lines are where event boundaries were used in fMRI analysis.

## Results

### Activation patterns in the posterior medial network, medial temporal network, and anterior temporal network contain event-specific representations at encoding and retrieval

Previous studies that used across-subject pattern similarity analysis (Chen et al., 2017) have shown that a number of neocortical areas carry information about events within movies. Here, we sought to identify event representations in the neocortical networks that interact most closely with the hippocampus. We focused these analyses the MTN, PMN, ATN and MPN (Figure 3A), based on previous work showing that these networks show high functional connectivity with the hippocampus (Barnett et al., 2021; see also Braga and Buckner, 2017 and Gordon et al., 2020).

**Figure 2.**
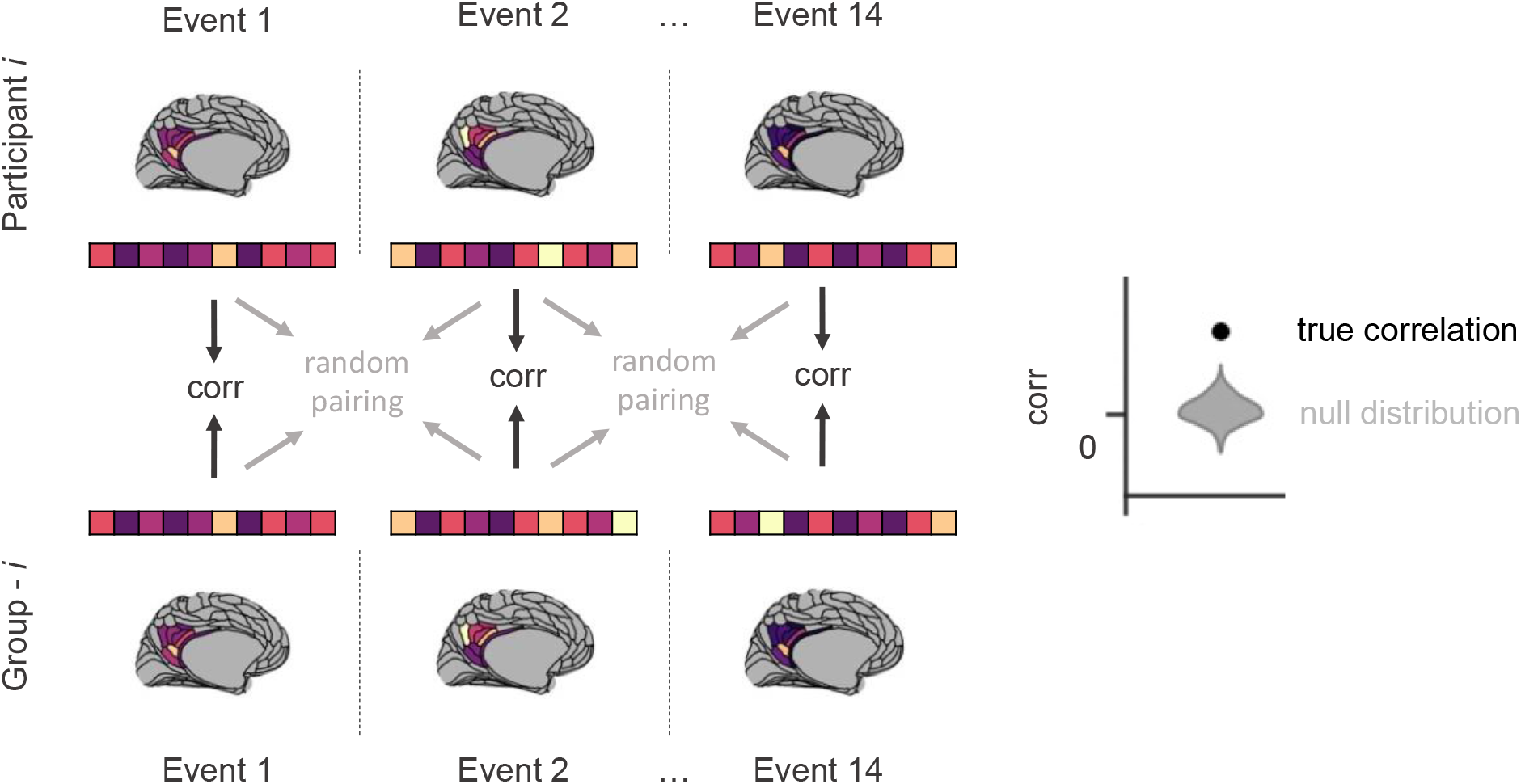
A visual schematic depicting the inter-subject pattern similarity (ISPS) analysis. For a given Participant *i* (top row), network level patterns of ROI activity are calculated for each event for each movie (depicted here are example data from the PMN on an inflated brain). The group averaged pattern excluding Participant *i* (bottom row) is also calculated. The vector of ROI patterns for the network (depicted by colored boxes) are correlated between the participant and group-averaged patterns (black arrows in the middle). To create a null distribution, this process is repeated 1000 times after randomly shuffling the event labels (grey arrows). A Schematic example of the mean of the true correlation between the participants and the group (the ISPS) which can be compared to the null distributions to determine significance is shown on the right.

**Figure 3.**
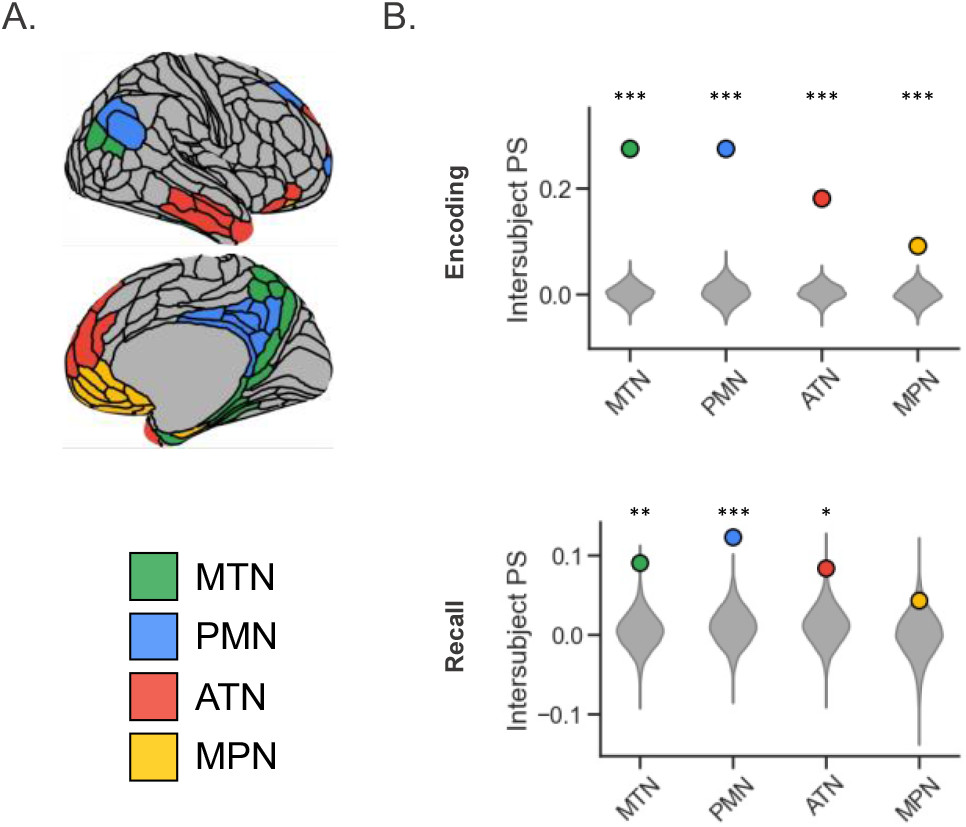
A. Regions of interest from the HCP-MMP atlas, color-coded by the cortico-hippocampal network to which they belong as identified in (Barnett et al., 2021). B. The average intersubject pattern similarity (PS) for each network represented by the circle and the grey violin plots display the group null distributions for each network. ATN, anterior temporal network; MPN, medial prefrontal network; MTN, medial temporal network; PMN, posterior medial network.

Events were defined as the time in between event boundaries that were designed into the stimuli and confirmed by an independent sample of 62 participants online (Figure 1B). The mean activity of each ROI in a network was calculated for each event, for each subject, resulting in one “multi-ROI activity pattern” per event, per subject, per network (see Figure 2 for an overview). This novel approach allowed us to assess the event representation of whole networks by using a multi-ROI activity pattern rather than the multi-voxel activity pattern within an ROI. The multi-ROI activity pattern of a network for each participant *i* was correlated with the averaged multi-ROI activity pattern for the rest of the group (excluding subject *i*) for each matching event and Fisher z-transformed to determine whether event-specific activation patterns were shared across participants. Significance was assessed via permutation testing which compared the average correlation to a null distribution (see Methods).

During encoding, this analysis revealed that shared event-specific patterns in the PMN (z = 13.7, p < .000001), MTN (z = 16.8, p < .0000001), ATN (z = 11.7, p < .000001), and MPN (z = 5.3, p < .000001). During recall, when participants had to internally generate event information, event-specific activation patterns were shared in the PMN (z = 4.8, p <.001), MTN (z = 3.8, p < .001), and the ATN (z = 2.9, p = .003), but not the MPN (z = 1.5, p = .13). Thus, shared event-specific representations were present in the PMN, MTN and ATN at both encoding and retrieval (Figure 3). These findings dovetail with previous work and suggest that these cortical networks carry event information that may be encoded into memory (Chen et al., 2017).

### Hippocampal to posterior medial network functional connectivity at event offset relates to subsequent retrieval success for events

Having established that event-specific activity patterns are present in the PMN, MTN, and ATN, we next tested how functional connectivity of these networks with the hippocampus relates to subsequent retrieval success for these events and whether this connectivity is particularly important at the offset of an event, as predicted by Lu, Hasson, & Norman (2022). For each subject and each event, we calculated the functional connectivity between the hippocampus and each ROI within the PMN, MTN, and ATN. Since the long-axis of the hippocampus has differential functional connectivity and specialization, we performed this separately for the anterior and posterior hippocampus (Brunec et al., 2017; Poppenk et al., 2013; Strange et al., 2014). These values were calculated at 20 TR (24.4s) time windows around each event’s onset, middle, and offset (see Methods). We averaged the hippocampal-ROI FC within networks to get a measure of hippocampal to cortical network functional connectivity (Figure 4A). The hippocampal-cortical FC for each network was entered as a regressor to predict subsequent recall using a generalized linear mixed-effects model. Memory for an event was determined by examining the free recall of each participant. If a participant recalled an occurrence from an event, then it was classified as recalled in a binary fashion. In the model we tested how delay (immediate vs. 2-day), hippocampal long-axis (anterior vs. posterior), and temporal window (start vs. middle vs. offset) interacted with the relationship between FC and memory by including these variables as regressors. We also included hippocampal boundary activity as a regressor to control for univariate activity effects (see Methods), since univariate activity in the hippocampus at stimulus offset has previously been shown to predict subsequent memory (Ben-Yakov and Dudai, 2011).

**Figure 4.**
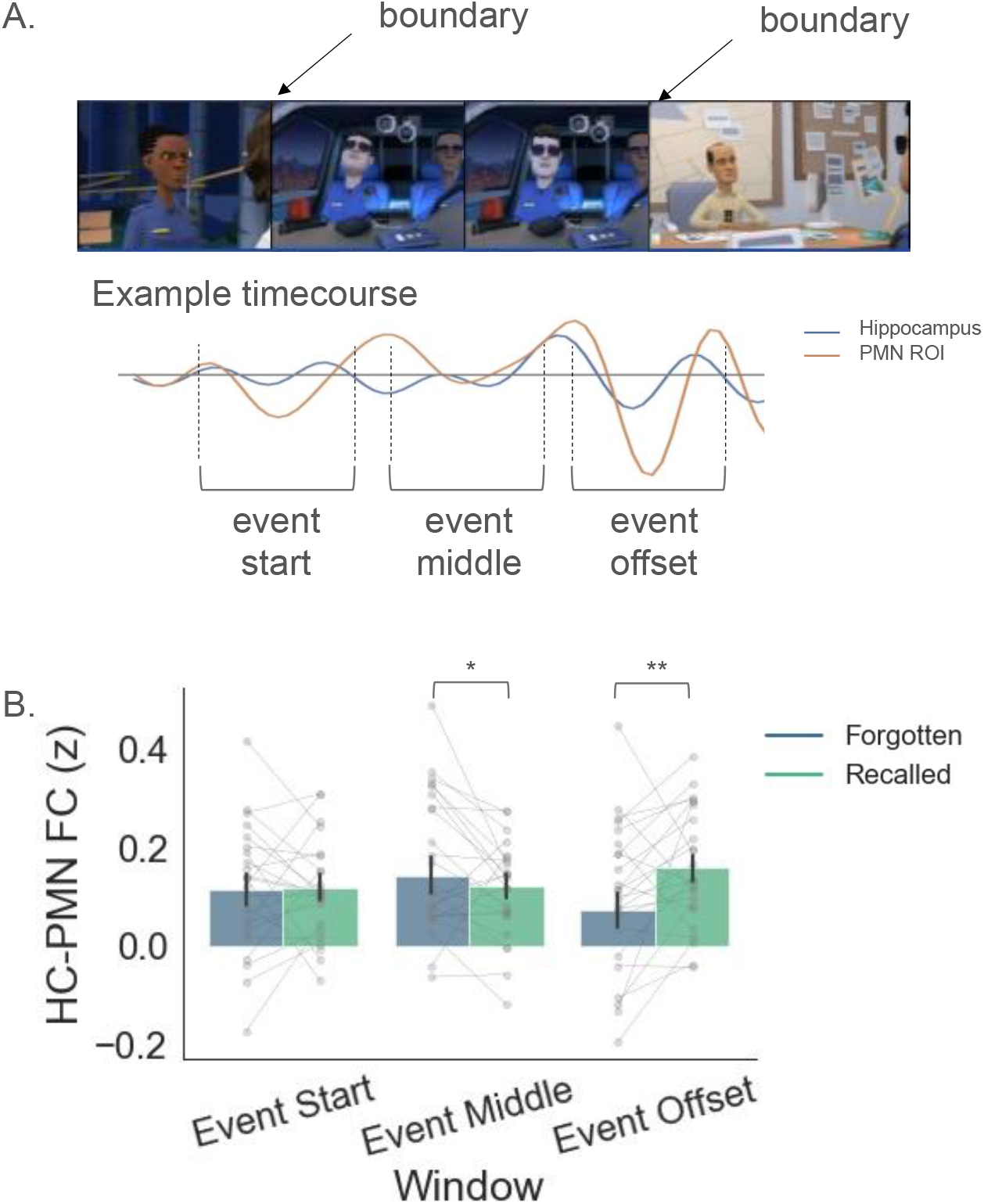
A. Schematic example of windows used for FC analysis. Ten TRs on either side of event boundaries, or the event middle point were used to estimate FC between the hippocampus and PMN. B. FC between the hippocampus and PMN at the event start, event middle, and event offset, split based on whether the event was subsequently recalled or not. Each small point represents a participant’s mean Hippocampal-PMN FC and each large dot with error bars represents the group mean and 95% confidence interval of the sample.

For the PMN, our analyses revealed a significant interaction between FC and window (Wald*X*^2^(2, N = 24) = 13.22, p = .001), but no interaction of FC with delay, hippocampal long-axis, univariate boundary activity or any combination of those factors (all p > .1). The significant interactions observed in this analysis reflected the fact that FC was differentially predictive of subsequent memory across different time windows, with significant effect at the event middle and event offset (Figure 4). At the event start, FC was not significantly related to subsequent recall, but during the middle window, we observed a significant effect such that higher FC was predictive of subsequent recall failure (z = 2.12, p = .044 FDR-corrected). To determine the consistency of this effect within the ROIs of this network, we repeated this analysis for each ROI in the PMN individually, finding that the estimated slope showed a negative relationship with memory in 21/24 ROIs suggesting that nearly all ROIs in the network shared this effect (Supplemental Figure 1).

Conversely, at the event offset, increased FC between the hippocampus and PMN was predictive of subsequent recall success (z = 3.08, p = .006, FDR-corrected). To determine the consistency of this effect, we repeated this analysis for each ROI in the PMN individually, finding that the estimated slope was positive in 24/24 ROIs demonstrating a consistent pattern of effects across the network (Supplemental Figure 1). These results indicate that the event offset is a critical moment for event encoding processes.

For the MTN, we observed a significant functional connectivity by window by delay interaction (*X*^2^(2, N = 24) = 11.5, p = .003). Follow-up on this interaction showed that higher functional connectivity between the hippocampus and MTN at the middle of an event during encoding, was associated with failure to recall that event at an immediate delay (z = 3.6, p = .001, FDR-corrected). To determine the consistency of this effect, we repeated this analysis for each ROI in the MTN individually, finding that the estimated slope was consistently negative in 28/30 ROIs suggesting that nearly all ROIs in the network shared this pattern of effects (Supplemental Figure 2). No other relationship between FC and memory was found at any other delays or windows (all p > .05, FDR).

For the ATN, we did not observe any significant relationship between functional connectivity and memory, nor any interactions with functional connectivity (all p > .05).

Across all models, we found that univariate hippocampal boundary activity was uniquely associated with subsequent recall success (*X*^2^ (1, N = 24) > 4.2, all p < .04), and this interacted with delay (*X*^2^ (2, N = 24) > 6.1, all p < .05), such that this effect was predictive at a 2-day delay (z > 3.2, all p < .003, FDR-corrected), but not at the immediate delay (z < .26, all p > .79, FDR-corrected) suggesting that the hippocampal boundary activity is important for forming stable memory representations. These results relate to the previous findings that hippocampal activity at the offset of a stimulus is associated with subsequent memory (Ben-Yakov and Dudai, 2011; Ben-Yakov et al., 2013). This effect is independent of the functional connectivity effects, as they were included in the same regression model.

### Hippocampal to posterior medial network functional connectivity at event offset relates to number of subsequently retrieved details

Having observed that increased FC between the hippocampus and PMN was predictive of subsequent memory success, we then asked whether hippocampal-PMN FC at encoding related to the number of details participants reported from the events that were successfully recalled. The audio recordings of each participants recall were transcribed and scored for the number of details by trained raters (JS & RY) using the scoring methods from the autobiographical interview (Levine et al., 2002). If hippocampal-PMN FC is a measure of successful encoding, then we should expect that events with higher FC will be recalled with more detail than those with low FC.

A mixed linear model to predict the number of details of a retrieved event showed a significant interaction between hippocampal-PMN FC, delay, and window (F(2, 2684.7) = 8.6, p = .0002). No other interactions of effects of interest were significant (all p > .05).

When breaking down the significant interactions, we observed that hippocampal-PMN FC at the event offset was positively associated with the number of details retrieved from an event at the 2-day retrieval phase (t(2684) = 3.2, p = .003, FDR-corrected; Figure 5), but not at the immediate recall phase (t(2684)= −1.1, p = .27, FDR-corrected). This finding dovetails with the above analysis and suggests that the coordinated activity between the hippocampus and PMN is particularly important for forming detailed, retrievable memories. To determine the consistency of this effect, we repeated this analysis for each ROI in the PMN individually, finding that the estimated slope showed this pattern in 22/24 ROIs suggesting that nearly all ROIs in the network shared this pattern of effects (Supplemental Figure 3). We also observed a trending effect that showed that hippocampal-PMN FC at the event onset was associated with the number of details retrieved from an event at immediate recall (t(2684) = 2.2, p = .06, FDR-corrected). Hippocampal-PMN FC at other windows and delays were not associated with the number of details retrieved for an event (all p > .09),

**Figure 5.**
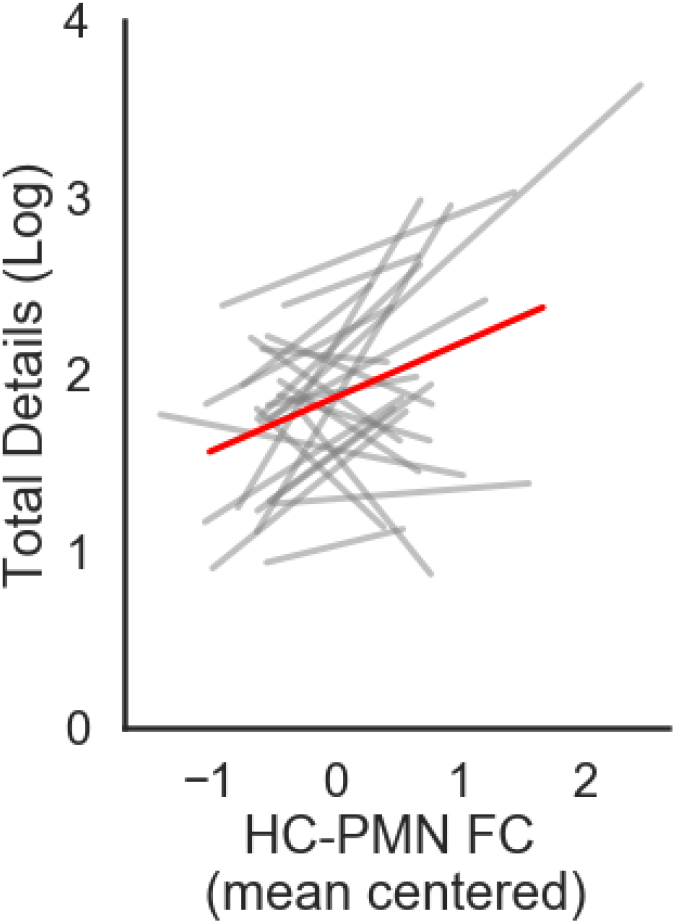
Relationship between HC-PMN FC at the event offset and the number of subsequently recalled details (log-transformed) for events that were successfully recalled at the 2-day delay. Each grey line represents the regression line of an individual participant, and each red line represents the group regression line produced from the linear mixed model.

## Discussion

Using dynamic, naturalistic stimuli, we demonstrated that interactions between the hippocampus and neocortical networks that carry event representations are critical for stable encoding. To do so, we first demonstrated that cortico-hippocampal networks (the PMN, MTN and ATN) carry event-specific information that is shared across participants during encoding and recall. Next, we showed that functional connectivity between the hippocampus and the PMN at event encoding was associated with subsequent recall success of the event. However, the timing of the interactions had a significant impact on this relationship such that hippocampal-PMN functional connectivity was associated with recall success at the event offset, but was associated with reduced likelihood of subsequent retrieval at the event middle. Further, the amount of detail retrieved after a delay was influenced by hippocampal-PMN interactions at encoding and, again, this relationship was modulated by the timing of the interactions such that higher functional connectivity between the hippocampus and PMN at the offset of an event was associated with more retrieved details. Our findings suggest that memory formation depends on communication between the hippocampus and neocortex at event boundaries, not for the event that is about to begin, but for the event that just finished.

Theories of episodic memory posit that event information represented in the neocortex is translated into a memory trace in the hippocampus (McClelland et al., 1995; Nadel and Moscovitch, 1997; Squire and Zola-Morgan, 1991; Teyler and DiScenna, 1986). The term “neocortex” is remarkably broad in these models, as much of the brain is important to our ongoing experience, but there is good reason to believe that a subset of the neocortex, the DMN, plays a specialized role, particularly for event memory. The DMN is functionally connected to the hippocampus (Barnett et al., 2021; Kahn et al., 2008) and has been discussed in terms of memory processing and retrieval (Ranganath and Ritchey, 2012; Robin and Moscovitch, 2017; Rugg and Vilberg, 2013), as it is known to be especially active during retrieval of autobiographical memories and recollection. As noted above, recent work has shown that the DMN can be subdivided into subnetworks (Barnett et al., 2021; Braga et al., 2019; Gordon et al., 2020; Ritchey and Cooper, 2020) and that these regions also play a role in encoding of naturalistic events (Baldassano et al., 2017; Chen et al., 2017; Oedekoven et al., 2017; Reagh and Ranganath, 2021). For example, Chen et al (2017) had participants encode a continuous, 50-minute segment of a television episode during fMRI scanning. They found that participants had shared event-specific multivariate patterns during both encoding and retrieval of events in DMN regions, particularly in posterior medial, and lateral parietal cortex—regions that are part of the PMN. These analyses were performed using the voxel-patterns within searchlight spheres across the brain. Here, we adopted a different approach to identify distributed event representations by characterizing the multi-ROI activation pattern with networks. We found shared, event-specific multi-ROI activation patterns across participants in the PMN, MTN and ATN. Given the event content represented in these regions, these neocortical regions may hold the event information that is ultimately encoded in hippocampal memory traces (Franklin et al., 2020; O’Reilly et al., 2021; Reagh and Ranganath, 2018).

Among the regions that showed event-specific patterns, our findings demonstrated that functional connections between the hippocampus and the PMN supported subsequent memory for events. This finding is consistent with the idea that event model features represented in the PMN are encoded in the hippocampus (Franklin et al., 2020). One recent computational modeling study has suggested that it might be optimal to selectively encode episodic information at the event offset, particularly when that memory will be subsequently used for comprehension and prediction (Lu et al., 2022). This model had a component that tracked features of the ongoing event, akin to event models represented in the PMN. The model also had a component for long term storage, like the hippocampus, and a dynamic gate that would allow information from the ongoing event model to be encoded into long-term storage. Opening that gate at the event offset—allowing for encoding at this moment—was ideal for model performance (Lu et al., 2022) which dovetails with our findings that PMN-hippocampal FC at the event offset was associated with higher retrieval success and, at the 2-day delay, more detailed retrieval of the events. Thus, the event offset may represent a window of time for features present in the ongoing event model to be encoded into accessible long-term memory representations, as was the case in the model proposed by Lu et al., (2022). In rodents and primates, detailed electrophysiological studies have also supported the idea that neocortical input into the hippocampus is gated via the rhinal cortex such that not all neocortical input that reaches the rhinal cortex is subsequently passed to the hippocampus (De Curtis and Paré, 2004). Additionally, the model proposed by Lu et al., (2022) also found that encoding in the middle of an event was detrimental to model performance which dovetails with our findings that higher FC between the hippocampus and PMN during the event middle was negatively associated with recall success. Functional interactions at the event middle between the hippocampus and the PMN and MTN may reflect the hippocampus encoding incomplete event information, it may reflect the hippocampus reinstating information into these networks, or it may reflect a reset of the event models which could negatively impact comprehension and retrievability of the rest of the event. While the data here cannot speak to the specific processing occurring, it does suggest that coordinated activity of the hippocampus and PMN during encoding is beneficial to retrieval specifically when this coordination occurs at the event offset.

An alternative account of event encoding has hypothesized that event models are represented in the hippocampus instead of the PMN (Griffiths and Fuentemilla, 2020; Milivojevic et al., 2016). For example, according to the model outlined by Griffiths and Fuentemilla (2020), as features of an event are encountered, unique populations of hippocampal neurons are activated to represent those features. The authors claim that this allows the hippocampus to build an ongoing event model of an event that is steadily and proactively encoded as it unfolds. At an event boundary the sequence of features is rapidly replayed to facilitate retroactive encoding of the final event representation. This logic suggests that ongoing interactions between the hippocampus and neocortex should be necessary and important for encoding, but makes no strong predictions about these interactions at the boundary. These predictions do not fit with our observations, as we found that hippocampal-cortical functional connectivity at the beginning and middle of an event were unrelated to memory (or related to worse memory), and only processing at the end related to subsequent retrieval success. There are several possible explanations that can reconcile these accounts. First, the model proposed by Griffiths and Fuentemilla (2018) is largely influenced by studies in which an event is described as a period in which a sequence of arbitrary stimuli is encoded under a stable context (Axmacher et al., 2010; Bahramisharif et al., 2018; Heusser et al., 2016). Because participants cannot form a predictive model of what to expect in these sequences based on any prior event schemas, the hippocampus may play a much stronger role in binding this sequence of arbitrary stimuli and the impact of posterior medial representations is low. When the sequence of events conforms to a familiar structure or schema, like the sequence of events when one visits an airport or restaurant, the PMN may have a more significant role in forming an event model with a limited role for the hippocampus as was shown in Baldassano et al., (2018). Further, when a sequence of information can be reliably predicted from a PMN event model, the hippocampus may not need to encode each encountered feature, but rather can simply index the PMN model which can reconstruct the elements. For example, people with hippocampal damage retain immediate recall of prose information (Baddeley and Wilson, 2002), can tell globally coherent stories (Keven et al., 2018), and even show similar activation patterns in posterior medial and lateral parietal cortex during movie viewing when compared to controls (Oedekoven et al., 2019; Zuo et al., 2020). This suggests that extrahippocampal regions such as the posterior medial cortex represent event models, especially when prior knowledge of events can inform episodes and memories. A modified version of Griffiths and Fuentemilla’s (2018) account could potentially fit our findings. It may be that, when encoding is particularly successful, the rapid replaying of features represented in the hippocampus at event boundaries leads to increased functional connectivity between the hippocampus and PMN. This could be true if the rapid replay leads to a rapid reinstatement of event features in neocortex. Given the temporal and spatial resolution limitations of fMRI, we are not able to examine rapid neuronal firing rates within a theta cycle in the hippocampus to determine if event features are incorporated into a sequence of population firings. However, using EEG Silva et al (2019) has shown that event boundaries trigger rapid replaying of multivariate patterns present in the just completed event (Silva et al., 2019). Thus, future studies using imaging techniques with higher temporal resolution should investigate whether the relationship between memory and hippocampal-PMN interactions at event boundaries is related to the rapid replay of hippocampal sequences that reinstate PMN representations.

The findings presented here seem to demonstrate a privileged role of the PMN in event memory, as we did not observe robust memory effects in other cortico-hippocampal networks. When examining multivariate event patterns, we did not find that the MPN showed consistent event-specific patterns during recall. Episodic memory studies have highlighted medial prefrontal cortex (mPFC) – hippocampal interactions as a major contributor to schema-related memory (Audrain and McAndrews, 2022; Ghosh and Gilboa, 2014; van Kesteren et al., 2012; Preston and Eichenbaum, 2013) and research using naturalistic stimuli has shown that multivariate patterns in the mPFC generalize between videos that share similar situations (Baldassano et al., 2018; Reagh and Ranganath, 2021) during viewing. The movie stimuli used in this study depicted narratives that tended to follow classic tropes of either a crime show or a fantasy show. Thus, MPN representations may have pertained to broader fantasy show vs. crime show schemas which could not differentiate between events within a movie.

While the MTN and ATN did show event-specific representations, their functional connectivity with the hippocampus was not associated with subsequent memory retrieval. Previous work has shown that the MTN may play a special role memory precision and perception (Koen et al., 2017; Kolarik et al., 2018; Ritchey and Cooper, 2020) and the ATN plays a special role in object, person, and social processing (DiNicola et al., 2020; Ranganath and Ritchey, 2012; Reagh and Ranganath, 2021). Memory assessment in this study was examined as overt free recall of the movies and did not impose any specific focus on precision, social elements, or conceptual elements of memory as participants were encouraged to retrieve every detail they could possibly remember. A memory task focussed more on the perceptual or social elements may be more related to hippocampal-MTN or hippocampal-ATN functional connectivity, respectively, during encoding and could be the subject of future research.

PMN event representations, in contrast, may be more generalized than MTN and ATN representations. For example, Zadbood et al. (2017) scanned participants as they both viewed narrative movies verbally recall those movies. They then played the verbal recall of the initial participants to a new set of participants during scanning. Remarkably, multivariate representations in the PCC and angular gyrus were shared in the brains of people watching the movies, those same people recalling the movies, and the new people listening to the recall of the movies (Zadbood et al., 2017) suggesting that these areas carry modality-invariant event representations. Even during episodic simulation, when participants must generate an event based on a cue, activation in the PCC and angular exclusively correlates with the number of generated details despite the fact that medial and anterior temporal regions are active for this type of task (Thakral et al., 2020). This broad involvement in event-based processing suggests a privileged role of the PMN in maintaining internal event representations necessary for encoding and reconstructing complex naturalistic memory.

Our study demonstrated that the influence of hippocampal-PMN functional connectivity on memory was dynamic. However, fMRI requires a relatively large window size for estimating functional connectivity (Hutchison et al., 2013) which, combined with low temporal resolution, limits the reliably to infer directionality of information transfer. Although we have hypothesized that event features represented in the PMN are bound by the hippocampus, we cannot conclude that the functional connectivity effects are driven by PMN to hippocampal information transfer. Further, given the time window used here to calculate functional connectivity, we may be capturing a back-and-forth conversation between the hippocampus and multiple brain regions of the PMN or MTN, rather than a one-time information packet transfer. However, one study that used intracranial recordings found that, at event boundaries during story listening, information flows from the cortex to the hippocampus which is what we would predict if this time point is important for encoding (Michelmann et al., 2021). Finally, we grouped regions together into subnetworks based on a group-averaged community detection solution from an independent sample of participants scanned at rest (Barnett et al., 2021). While we were able to find robust effects, recent work has shown that there are individual differences in network organization across participants (Braga et al., 2019; Gordon et al., 2017a). These individual network estimations are mostly overlapping with group estimated networks (Gordon et al., 2017b), but future work may be able to rely on individually estimated networks to characterize subtler memory effects.

In conclusion, the present study has presented evidence to extend our understanding of how the brain translates experience into memory by examining the timecourse of interactions between the hippocampus and neocortex. We discovered that interactions between hippocampus and PMN at the offset of an event, but not the onset or middle, is beneficial for subsequent recall and, higher functional connectivity at the offset is associated with more detailed retrieval when events are retrieved after a longer delay. This suggests that the event offset is a critical moment in time for encoding complex, narrative events.

## Methods

### Participants

For the primary resting-state fMRI dataset, twenty-nine healthy, young adult participants were recruited from the University of California, Davis, and surrounding area. Five participants were excluded due to scanner artifact or audio recording failures during recall leaving a final sample size of twenty-four (N_Females_ = 14, mean age = 23.5 years [SD = 4.1 years]). All participants were right-handed and neurologically healthy. The study was approved by the Institutional Review Board of the University of California at Davis (IRB #1352490 & IRB # 637028) and adheres to all principles of The Belmont Report. All participants provided written informed consent prior to participation. Participants were compensated $20/hour for their time.

### Experimental Stimuli

We constructed a set of animated, short movies using Plotagon (https://www.plotagon.com/desktop/). All movies were scripted and produced by the first author (AJB) with voice acting provided by volunteers in the center and with assistance from author (BICS) for audio preprocessing in Audacity v2.4.2. All movies depicted a continuous narrative. A practice movie was created that depicted two characters in conversation and was one-minute in length. The two experimental movies used in scanner were approximately 16.5 minutes in length each (Movie 1, 16:47; Movie 2, 16:22). Both movies consisted of a series of events that each depicted two characters at a time interacting. Each had a total of seven unique characters, and each had a total of eight spatial locations. One movie was set in a medieval, fantasy-like world and the other depicted a crime scene investigation narrative. The movies were designed such that each would elicit 13 event boundaries corresponding to changes in location, changes in time, or changes in the combination of characters shown.

### Event boundary rating

An independent sample of 97 online participants viewed one of the two stimulus movies via the online platform Testable (www.testable.org). Participants were presented with one of the two movies, and during viewing, were asked to press the spacebar whenever they perceived that a meaningful event had ended, and another event had begun. Online participant response data was examined for data quality and participants who made no response (29 participants) and those that responded outside the normal distribution of the sample (6 subjects) were excluded resulting in a sample of 62 participants. Summed participant inputs over time were modelled with a kernel density function with a 5s bandwidth for each movie and confirmed the temporal location of the intended event boundaries (Figure 1B). One unplanned, but reliable, event boundary was observed in both movies. One unplanned event boundary pertained to characters changing positions within the same scene as they continued the same conversation (characters moved from standing at a bar to sitting at a table). The other unplanned event pertained to a fade to a black screen with written text describing a change in location. The periods in between event boundaries were considered events with each movie having a total of 14 events.

### Pre-scan Procedures

#### Practice Recall

Prior to scanning, participants completed a brief example trial of the experiment to get accustomed to the type of task they would be performing in the scanner. After providing written consent, participants viewed a one-minute cartoon video on a desktop personal computer. They were instructed to “watch the video as you would a television show you are interested in” and that they would be asked to remember the video after. They were asked to recall, in as much detail as possible everything they could from the video. They were also asked to recall in temporal order, if possible, but completeness and detail were considered more important than temporal order and, so, if at any point they realized they had missed something, they were instructed to return to that detail. They were instructed to mention every detail they could recall, even if it seemed irrelevant. After participants performed this recall, the experimenter provided a general probe to the participant asking if there was anything else they could recall. Recall data was not recorded, but participants were encouraged to push themselves to retrieve all details they could once they performed the task in the scanner.

#### Character Familiarization

After responding to the practice video, participants were then familiarized with a set of characters that they would subsequently view in upcoming movie stimuli in the MRI scanner task. Participants were presented with a total of 14 character-avatars using PsychoPy v3.0.0 (Peirce et al., 2019). Participants were told that they would see a series of cartoon people with a name and a fact about each character and they were instructed to try to memorize the names. The 14 characters were split into two blocks of seven characters, based on the two movies in which they would subsequently feature. Each character avatar was presented one at a time, for 3000ms, in the center of the computer screen and the character’s name was printed underneath the character’s avatar. In between each character presentation was a 1000ms interstimulus interval. Following encoding of each character and name, participants performed a two-alternative forced-choice decision task for each of the character’s names. Participants were shown each character avatar individually, in random order, with two names (one originally shown with the displayed character, and one from another character that was previously shown) underneath the character. Participants were instructed to indicate which name belonged to the displayed character. A 1000ms inter trial interval was present between each testing trial. After this testing block, participants were once again shown each character avatar, with their name printed underneath, along with a fact about that character’s role in the upcoming videos. For example, the character named “Arlene” was presented underneath the cartoon avatar of the Arlene character alongside Arlene’s role in the video of “Barmaid”. This provided feedback on the names for the participant and added an additional semantic fact for the participants to encourage memory formation for the characters.

#### Pre-scan set-up

Participants were fitted with MRI-compatible earbuds with replaceable foam inserts (https://www.sens.com/products/model-s14/) and were provided with additional foam padding inside the head coil for hearing protection and to mitigate head motion. Participants were additionally given bodily padding, blankets, and corrective lenses as needed. An MRI compatible microphone (Optoacoustics FOMRI-III; https://www.optoacoustics.com/medical/fomri-iii) with bidirectional noise-cancelling was affixed to the head coil, and the receiver (covered by a disposable sanitary pop screen) was positioned over the participant’s mouth. Participants were given a description of strategies to remain still while speaking during functional image acquisition. Participants were told to avoid nodding or shaking their head and have minimal jaw movement—by keeping their teeth touching—during speaking. During MRI data acquisition, an eye tracker was operational to monitor participants’ wakefulness and head motion during spoken recall, but no eye tracking data were recorded.

#### fMRI task overview

Participants were tasked with encoding and recalling two, 16.5-minute animated videos. One video was recalled after a brief-minute delay, and the other was recalled after a 48-hour delay. Participants were split into four groups to counterbalance video order and retention delay order—half of the participants performed the short delay recall first, and half performed the 48-hour delay recall first (Figure 1A).

Encoding block instructions: Prior to each encoding scan, participants were instructed to: “Watch the following movie as you would a television show you are interested in. We will ask you to remember the movie at a later time.” Participants then viewed the first of the two animated videos. After encoding, participants underwent a 3.5-minute anatomical scan.

Recall block instructions: During recall scans they were instructed to: “In as much detail as possible tell me everything you can remember about the last movie we showed you. Try to recount the events in the original order in which they occurred. Completeness and detail are more important than temporal order. If at any point you realize that you have missed something, return to it. Try to describe EVERY detail that you have about the movie you just watched, even if it seems irrelevant.” The participants then spoke into the scanner safe microphone and recalled details from the video they had most recently viewed during a functional scan. Participants received additional instructions regarding head movement prior to recall: “Remember to stay as still as possible. Try to speak while moving only your lips and not your jaw; Sometimes people tend to nod or shake their head when they are talking. Avoid these tendencies as much as possible.”

If the video was scheduled for the short recall delay, the participant would perform the recall block immediately after the anatomical scan. If the video was in scheduled for the 48-hour delay first recall group, the participant would then undergo a reverse phase encoding functional scan, followed by a diffusion weighted scan, and a reverse phase encoded diffusion weighted scan (to be reported elsewhere). The participant would then leave the scanner and return two days later to complete the recall block in the scanner.

After completing the encoding and recall of both movies, participants then watched an 11-minute documentary-style video and were instructed to watch the following video as they would a television show they are interested in. This final video was not recalled by the participants.

#### Recall scoring

Audio recall from each participant’s fMRI recall run was transcribed automatically using the python tool SpeechRecognition (https://github.com/Uberi/speech_recognition#readme) and the automatic transcription was examined, and any transcription errors were corrected by two trained research assistants (J.S. and R.Y.). These two trained research assistants parsed the transcripts into details using an adapted version of the autobiographical interview (Levine et al., 2002). A detail “was defined as a unique occurrence, observation, or thought, typically expressed as a grammatical clause. Additional information in the clause was scored separately”. These details were then categorized as either central details, or peripheral details to determine whether details that are central to the narrative plot are better retained than those that are peripheral. Central and peripheral details were both details that could be verified as true from the information presented in the movies—the combination of central and peripheral details make up all total verifiable details (Cohn-Sheehy et al., 2021). To categorize central and peripheral details, four research assistants watched each video and created an annotation file for each line of dialogue and occurrence they viewed. This was then combined into a single document and the research assistants ranked each detail/occurrence on a scale of 1-5 in terms of detail centrality. Raters were instructed that a detail would be considered central if removal of that detail would affect their interpretation or comprehension of the narrative. A rating of 5 would be given if removal of the detail significantly affected their interpretation or comprehension and a 1 was to be given if removal of the detail would have no effect at all on their interpretation or comprehension of the narrative. Details that were given an average score greater than 3.5 were considered central details and summary statements of these details were produced via conferencing among the group. During scoring, details were classified as central if they matched a detail that was identified as central by the central detail rating group, and all other verifiable details were classified as peripheral. While this scoring provided a fine-grained approach to evaluating memory, we also examined whether events were recalled in a binary fashion. If any event-specific details were recalled from an event, that event was said to be successfully recalled and events for which no details were recalled were classified as forgotten or unsuccessfully recalled. Analysis of details can be found in the Supplemental Results.

#### MRI Acquisition

MRI scanning for the primary dataset was performed using a 3T Siemens Skyra scanner system with a 32-channel head coil. T1-weighted structural images were acquired using a magnetization prepared rapid acquisition gradient-echo pulse sequence (TR = 1900 ms; TE = 3.1 ms; field of view = 256 mm^2^; flip angle = 7°; image matrix = 256 × 256, 208 axial slices with 1.0 mm^3^ voxel size). Functional images were acquired using a gradient EPI sequence (TR = 1220 ms; TE = 24 ms; field of view = 192 mm^2^ ; image matrix = 64 × 64; flip angle = 67°; multiband factor = 2; 38 interleaved axial slices, voxel size = 3 mm^3^ isotropic), phase encoding: anterior-posterior. Reverse phase-encoded EPI data was also acquired using the same parameters to correct for phase encoding distortion in preprocessing (below).

### MRI preprocessing

#### Anatomical data preprocessing

A total of 2 T1-weighted (T1w) images were found within the input BIDS dataset. All of them were corrected for intensity non-uniformity (INU) with N4BiasFieldCorrection (Tustison et al., 2010), distributed with ANTs 2.3.3 (Avants et al., 2008) (RRID:SCR_004757). The T1w-reference was then skull-stripped with a Nipype implementation of the antsBrainExtraction.sh workflow (from ANTs), using OASIS30ANTs as target template. Brain tissue segmentation of cerebrospinal fluid (CSF), white-matter (WM) and gray-matter (GM) was performed on the brain-extracted T1w using fast (Zhang et al., 2001) (FSL 5.0.9, RRID:SCR_002823). A T1w-reference map was computed after registration of 2 T1w images (after INU-correction) using mri_robust_template (Reuter et al., 2010) (FreeSurfer 6.0.1). Brain surfaces were reconstructed using recon-all (Dale et al., 1999) (FreeSurfer 6.0.1, RRID:SCR_001847), and the brain mask estimated previously was refined with a custom variation of the method to reconcile ANTs-derived and FreeSurfer-derived segmentations of the cortical gray-matter of Mindboggle (Klein et al., 2017) (RRID:SCR_002438). Volume-based spatial normalization to one standard space (MNI152NLin2009cAsym) was performed through nonlinear registration with antsRegistration (ANTs 2.3.3), using brain-extracted versions of both T1w reference and the T1w template. The following template was selected for spatial normalization: ICBM 152 Nonlinear Asymmetrical template version 2009c (Fonov et al., 2009) (RRID:SCR_008796; TemplateFlow ID: MNI152NLin2009cAsym). Surface-based registration to the HCP-MMP1.0 atlas (Glasser et al., 2016) was performed, and subject-specific cortical regions of interest (ROIs) were calculated according to atlas boundaries. Surface-based cortical regions were converted to volumetric ROIs and transformed into functional native space. The hippocampus was segmented in FreeSurfer in an automated fashion. Manual adjustments were done to correct mis-classified voxels, and the hippocampus was divided into anterior and posterior segments based off the disappearance of the uncal apex (Poppenk et al., 2013), with the posterior hippocampus designated as all of the hippocampus posterior to the disappearance of the uncal apex on a coronal slice.

#### Functional data preprocessing

For each of the BOLD runs found per subject (across all tasks and sessions), the following preprocessing was performed. First, a reference volume and its skull-stripped version were generated using a custom methodology of fMRIPrep. A B0-nonuniformity map (or fieldmap) was estimated based on two (or more) echo-planar imaging (EPI) references with opposing phase-encoding directions, with 3dQwarp (Cox and Hyde, 1997) (AFNI 20160207). Based on the estimated susceptibility distortion, a corrected EPI (echo-planar imaging) reference was calculated for a more accurate co-registration with the anatomical reference. The BOLD reference was then co-registered to the T1w reference using bbregister (FreeSurfer) which implements boundary-based registration (Greve and Fischl, 2009). Co-registration was configured with six degrees of freedom. Head-motion parameters with respect to the BOLD reference (transformation matrices, and six corresponding rotation and translation parameters) are estimated before any spatiotemporal filtering using mcflirt (Jenkinson et al., 2002) (FSL 5.0.9). The BOLD time-series were resampled onto the following surfaces (FreeSurfer reconstruction nomenclature): fsaverage5. The BOLD time-series (including slice-timing correction when applied) were resampled onto their original, native space by applying a single, composite transform to correct for head-motion and susceptibility distortions. These resampled BOLD time-series will be referred to as preprocessed BOLD in original space, or just preprocessed BOLD. The BOLD time-series were resampled into standard space, generating a preprocessed BOLD run in MNI152NLin2009cAsym space. First, a reference volume and its skull-stripped version were generated using a custom methodology of fMRIPrep. Several confounding time-series were calculated based on the preprocessed BOLD: framewise displacement (FD), DVARS and three region-wise global signals. FD was computed using two formulations following (Power et al., 2014) (absolute sum of relative motions) and (Jenkinson et al., 2002) (relative root mean square displacement between affines). FD and DVARS are calculated for each functional run, both using their implementations in Nipype (following the definitions by (Power et al., 2014)). The three global signals are extracted within the CSF, the WM, and the whole-brain masks. The head-motion estimates calculated in the correction step were also placed within the corresponding confounds file. Frames that exceeded a threshold of 0.5 mm FD or 3.0 standardised DVARS were annotated as motion outliers. All resamplings can be performed with a single interpolation step by composing all the pertinent transformations (i.e. head-motion transform matrices, susceptibility distortion correction when available, and co-registrations to anatomical and output spaces). Gridded (volumetric) resamplings were performed using antsApplyTransforms (ANTs), configured with Lanczos interpolation to minimize the smoothing effects of other kernels (Lanczos, 1964). Non-gridded (surface) resamplings were performed using mri_vol2surf (FreeSurfer). Following preprocessing from fMRIprep, a set of nuisance regressors was made for each participant. Translational motion in the x, y, and z direction and rotational motion in pitch, roll, and yaw were calculated for each run for each subject. Scans with greater than 0.5 mm framewise displacement or DVARS > 3 were flagged as outlier TRs and a regressor was made for each outlier TR. Finally, the global signal of each TR was calculated. For the functional connectivity analysis, each of these nuisance regressors was entered into a model and regressed out from the fMRI timecourse of the participants and the timecourse was bandpass filtered to restrict frequencies to .009 —.08Hz using the tproject function from AFNI (Cox and Hyde, 1997) via nipype version 1.4.0 (Gorgolewski et al., 2011) to create a confound corrected timeseries for each participant, for each run. For the intersubject pattern similarity analysis, nuisance regressors were entered into a model and regressed out from the fMRI timecourse of the participants and the timecourse was high-pass filtered minimally to remove frequencies below .001Hz using the tproject function described above.

#### Intersubject pattern similarity analysis

Prior to addressing our primary hypotheses, we first confirmed that the cortico-hippocampal networks in our sample showed event specific representations. To do so, we adopted the intersubject representational similarity analysis used in (Chen et al., 2017). We focused these analyses the MTN, PMN, ATN and MPN (Figure 3A), based on previous work showing that these networks show high functional connectivity with the hippocampus (Barnett et al., 2021; see also Braga and Buckner, 2017 and Gordon et al., 2020) the MTN and three DMN subnetworks. The mean BOLD activity of each ROI in a network was calculated for each event, for each subject, resulting in one “multi-ROI activity pattern” per event, per subject, per network (see

Figure 2 for an overview). The multi-ROI activity pattern of a network for each participant *i* was correlated with the averaged multi-ROI activity pattern for the rest of the group (excluding subject *i*) for each matching event and Fisher z-transformed. The average Fisher transformed z-value for the matching events was calculated and compared to a null distribution that was calculated by repeating this procedure 5000 times, scrambling the event labels on each iteration. If a network carried event-specific information, then the average correlation of the matching events should be significantly higher than the null distribution of randomly correlated events. The values of the true correlations for the group were thus compared to the null distribution to assess significance. This was done for both movie encoding and recall, separately. For recall, for each subject *i*, we calculated the multi-ROI activity pattern for every event *e* that was recalled by subject *i* and correlated it to the average multi-ROI activity pattern for the rest of the group that also remembered event *e*, as was done in (Chen et al., 2017).

#### Functional connectivity analysis

Pairwise FC was estimated between every pair of ROIs in the brain by correlating the confound corrected timeseries between each pair of ROIs, for each subject, during each functional run. The resulting Pearson’s correlations were then Fisher z-transformed. To assess the FC between the hippocampus and a cortico-hippocampal networks of interest, we calculated the average FC of the hippocampus to all the ROIs in the cortico-hippocampal network. FC estimations were performed at three time-windows of interest—the event onset, middle and offset. For each event, we took a 20 TR window (24.4s) around the event onset, a 20 TR window around the event offset, and a 20 TR window around the event middle—a window length previously shown to be sensitive to BOLD signal correlations using naturalistic movie stimuli (Lin et al., 2019; Mukamel et al., 2005). This was done separately for anterior and posterior hippocampus, as some evidence suggests that there are different functions and different network connections along the hippocampal long-axis (Adnan et al., 2016; Poppenk and Moscovitch, 2011; Strange et al., 2014).

#### Univariate boundary activity

The anatomically registered functional data was entered into single subject general linear models for each subject and each movie using SPM12. A separate regressor was made for each event boundary and each event middle (the arithmetic middle of the event) having durations of 0s and were convolved with the canonical hemodynamic response function in SPM12. Nuisance regressors for motion (pitch, roll, yaw, x-translation, y-translation, z-translation) along with regressors flagging outlier TRs of high motion (FD > 0.5 or DVARS > 3) were entered as regressors of no interest. We also attempted to control for low-level visual information in our GLM by using the routine outlined by Reagh et al (2020). Edge pixels were calculated via a python routine, which read in each video stimulus, split it into its constituent frames, and in an automated fashion performed edge-detection on each frame (using python package opencv). The proportion of edge pixels to total pixel count was calculated (NumPy) for each frame, and was output into a comma-separated value file. We then resampled frame-wise edge information to correspond to the temporal scale of the fMRI data by averaging across adjacent frames within the interval of each TR (1220 ms). This resultant temporally smoothed vector served as our estimate of low-level visual information in each timepoint of the video. This visual information vector for each video was entered into the GLMs as a regressor of non-interest. Data underwent a high-pass filter with a cut-off of 128s.

For each subject GLM, for each movie, we estimated the beta values for each event boundary and event middle at every voxel in the brain. We extracted the beta values for these regressors averaged within the anterior and posterior hippocampus of each subject using the ROI extraction toolbox (Whitfield-Gabrieli and Nieto-Castanon, 2012). To calculate the univariate boundary effect in the hippocampus, we subtracted the beta value of the event middle from the beta value of the event boundary for each event.

#### Subsequent Memory effects analysis

The hippocampal to network FC values were mean centered within each participant for each cortico-hippocampal network of interest that showed event-specific multivariate patterns. These mean-centered values were entered into our statistical models. A generalized linear mixed model was used to run a logistic regression determining whether subsequent recall success could be predicted from FC, delay condition (immediate vs. two-day delay), the interaction of these two terms, and whether these interacted with FC window (beginning vs. middle vs. offset), and hippocampal long-axis (anterior vs. posterior) using glmer from the lme4 package in R. Since hippocampal boundary activity has previously been associated with recall success, we included boundary activity as a regressor in the model. Significance for terms in the logistic generalized linear mixed model was determined using Wald Chi-square tests. A random intercept was entered into the model for each participant. Movie label was entered as a covariate of no interest. Follow-up effects were calculated using emtrends from the emmeans package in R.

A second model was constructed using only events that were successfully recalled to determine the influence of FC on the amount of detail remembered from events. Since we observed no differences in central and peripheral details (Supplemental Results), here the dependent variable was the log-transformed total numbers of details recalled from events and the predictors were the same as above. Details were log-transformed since a Shapiro-Wilk test revealed that the detail distribution was not normal, W = .72, p < .001. This analysis was modelled using a linear mixed model with lmer from the lme4 package in R. Thus, rather than examining how FC relates to success of recall of events, we examined how it relates to the amount of detail retrieved when the event is recalled. Follow-up effects were calculated using emtrends from the emmeans package in R. Statistical probabilities were adjusted for multiple comparisons using the FDR method (Benjamini and Yekutieli, 2001) implemented in the emmeans package.

## Supporting information

Supplemental Methods and Results

## Supplemental Methods

### Recall statistical analysis

A mixed linear model was used to determine the effect of delay on the number of events subsequently recalled, and the number of details recalled. An event was considered recalled if participants mentioned any detail that was specific to a given event. Movie and counter-balance group were entered as covariates of no interest. The model predicting number of details recalled from delay also included detail type (central vs. peripheral) as an effect of interest. The Shapiro–Wilk test was used to evaluate normality on the distribution of details.

## Supplemental Results

### Recall statistical analysis

Participants retrieved a fewer number of events after a 2-day delay compared to the immediate recall condition (t(44) = 4.91, p < .001). The distribution of the number of details recalled by participants was skewed, with some subjects recalling many more details above average than the group. A Shapiro-Wilk test revealed that the distribution was not normal, W = .72, p < .001. The distribution was therefore log-transformed using the natural log, resulting in a normal distribution, W = 0.97, p = .058. Using the log-transformed detail scoring, we observed that participants reported fewer details after a 2-day delay compared to the immediate recall condition (t(88) = 3.2, p = .001). Participants also tended to report more peripheral details than central details (t(88) = 8.7, p < .001), but there was no interaction between detail type and delay in terms of details recalled (t(88) = .36, p = .7. In all models, there was no effect of movie, (t-values < 1, p-values > 0.29) or counter-balance group, (F-values < 1.76, p-values > .17). Since there were no interactions with detail type, future analysis pooled details into total details.

---

**Supplemental Figure 1.**
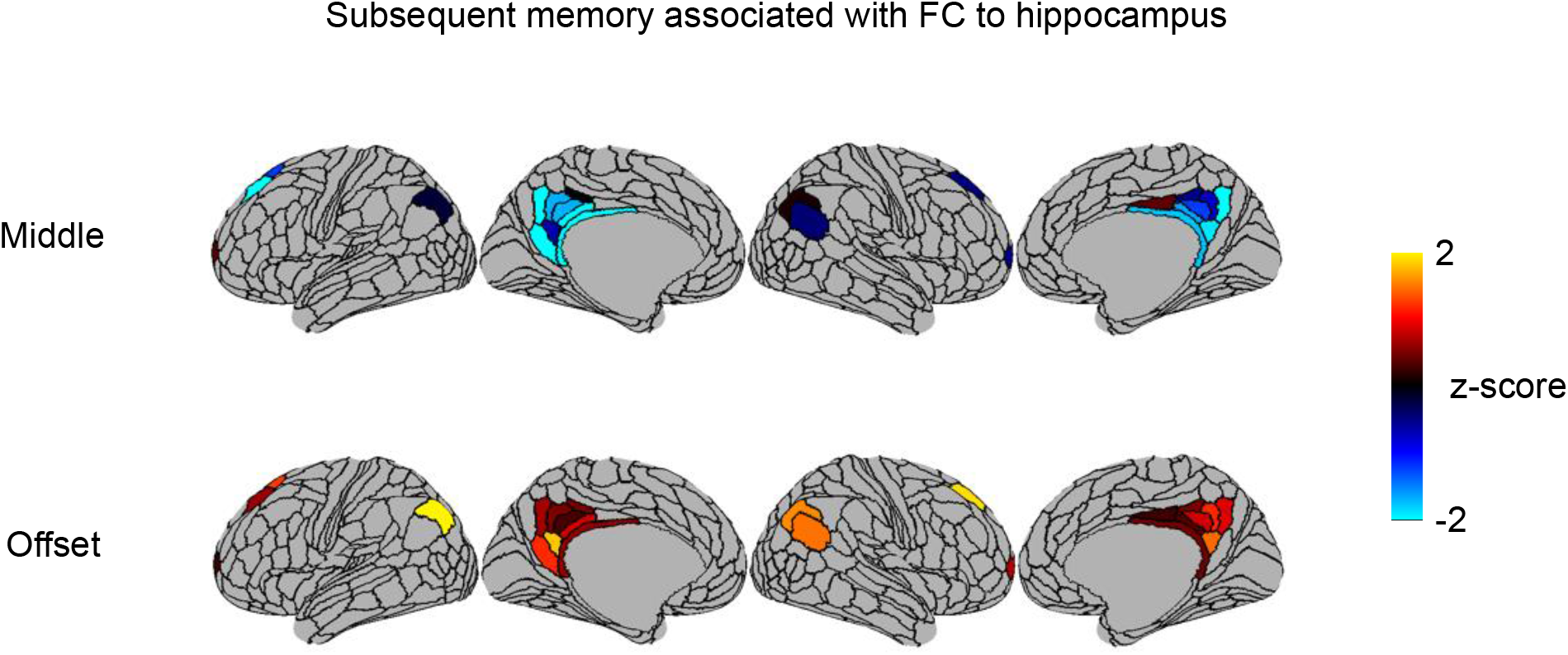
Relationship between subsequent recall success and hippocampal FC with ROIs in the PMN plotted on an inflated brain surface. Warm colors indicate that functional connectivity between the ROI and hippocampus is associated with better subsequent memory for events and cool colors indicate that functional connectivity with the hippocampus is associated with worse subsequent memory.

**Supplemental Figure 2.**
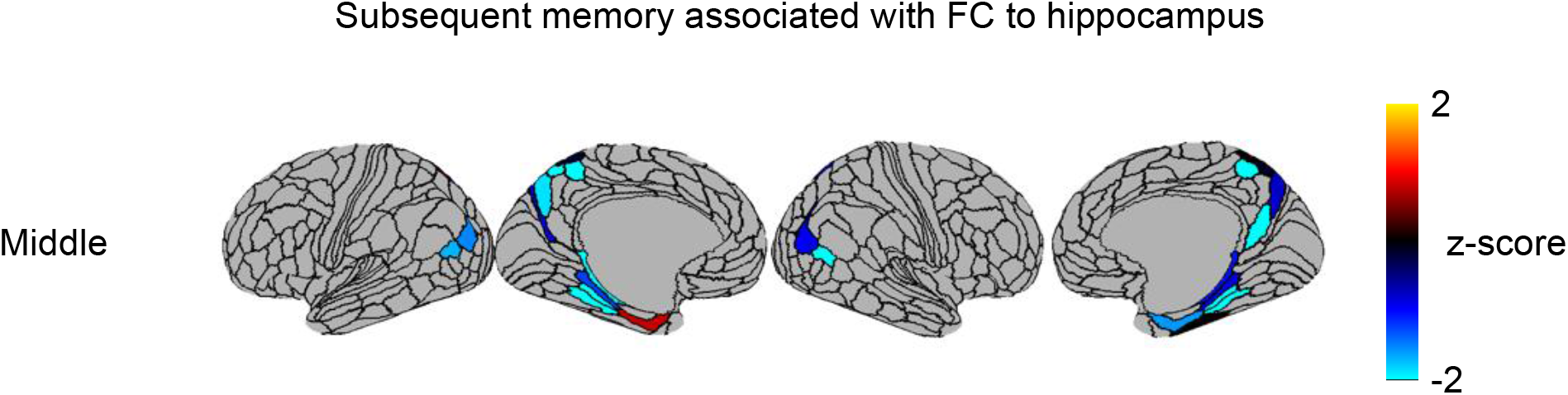
Relationship between subsequent recall success and hippocampal FC with ROIs in the MTN at the immediate recall time point plotted on an inflated brain surface.

**Supplemental Figure 3.**
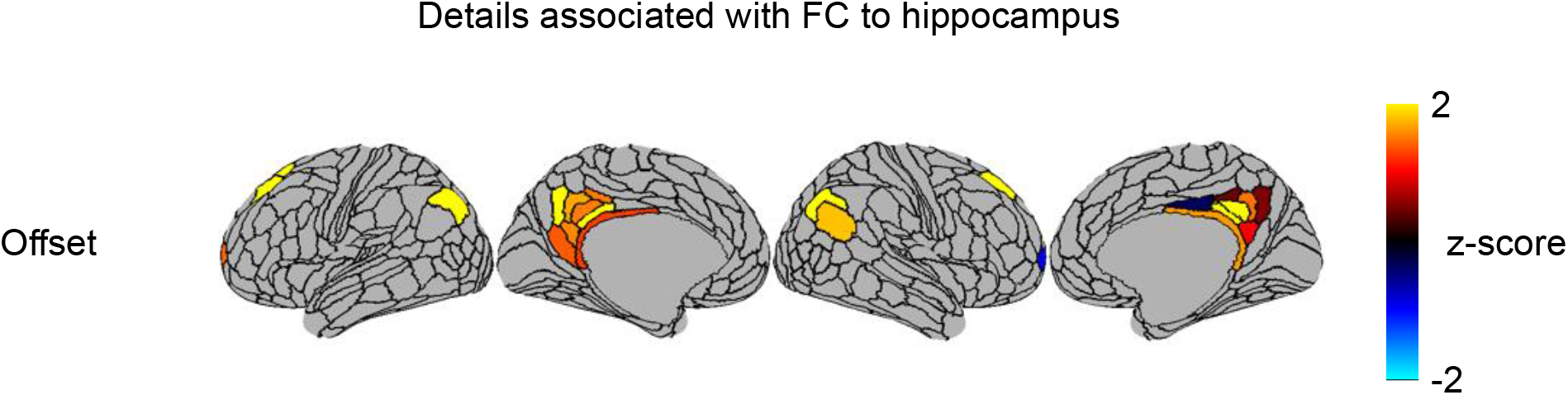
Relationship between number of subsequently recalled and hippocampal FC with ROIs in the PMN at the delayed recall time point plotted on an inflated brain surface.

## References

Adnan, A., Barnett, A., Moayedi, M., McCormick, C., Cohn, M., and McAndrews, M.P. (2016). Distinct hippocampal functional networks revealed by tractography-based parcellation. Brain Struct. Funct. 221, 2999–3012. https://doi.org/10.1007/s00429-015-1084-x.

Andrews-Hanna, J.R., Reidler, J.S., Sepulcre, J., Poulin, R., and Buckner, R.L. (2010). Functional-Anatomic Fractionation of the Brain’s Default Network. Neuron 65, 550–562. https://doi.org/10.1016/j.neuron.2010.02.005.

Audrain, S., and McAndrews, M.P. (2022). Schemas provide a scaffold for neocortical integration of new memories over time. Nat. Commun. 13, 5795. https://doi.org/10.1038/s41467-022-33517-0.

Avants, B., Epstein, C., Grossman, M., and Gee, J. (2008). Symmetric diffeomorphic image registration with cross-correlation: Evaluating automated labeling of elderly and neurodegenerative brain. Med. Image Anal. 12, 26–41. https://doi.org/10.1016/j.media.2007.06.004.

Axmacher, N., Henseler, M.M., Jensen, O., Weinreich, I., Elger, C.E., and Fell, J. (2010). Cross-frequency coupling supports multi-item working memory in the human hippocampus. Proc. Natl. Acad. Sci. U. S. A. 107, 3228–3233. https://doi.org/10.1073/pnas.0911531107.

Baddeley, A., and Wilson, B.A. (2002). Prose recall and amnesia: Implications for the structure of working memory. Neuropsychologia 40, 1737–1743. https://doi.org/10.1016/S0028-3932(01)00146-4.

Bahramisharif, A., Jensen, O., Jacobs, J., and Lisman, J. (2018). Serial representation of items during working memory maintenance at letter-selective cortical sites. PLOS Biol. 16, e2003805. https://doi.org/10.1371/journal.pbio.2003805.

Baldassano, C., Chen, J., Zadbood, A., Pillow, J.W., Hasson, U., and Norman, K.A. (2017). Discovering Event Structure in Continuous Narrative Perception and Memory. Neuron 95, 709–721.e5. https://doi.org/10.1016/j.neuron.2017.06.041.

Baldassano, C., Hasson, U., and Norman, K.A. (2018). Representation of Real-World Event Schemas during Narrative Perception. J. Neurosci. 38, 9689–9699. https://doi.org/10.1523/JNEUROSCI.0251-18.2018.

Barnett, A.J., Reilly, W., Dimsdale-Zucker, H.R., Mizrak, E., Reagh, Z., and Ranganath, C. (2021). Intrinsic connectivity reveals functionally distinct cortico-hippocampal networks in the human brain. PLOS Biol. 19, e3001275. https://doi.org/10.1371/journal.pbio.3001275.

Bartlett, F.C. (1932). Remembering: A Study in Experimental and Social Psychology.

Ben-Yakov, A., and Dudai, Y. (2011). Constructing realistic engrams: Poststimulus activity of hippocampus and dorsal striatum predicts subsequent episodic memory. J. Neurosci. 31, 9032–9042. https://doi.org/10.1523/JNEUROSCI.0702-11.2011.

Ben-Yakov, A., and Henson, R.N. (2018). The hippocampal film editor: Sensitivity and specificity to event boundaries in continuous experience. J. Neurosci. 38, 10057–10068. https://doi.org/10.1523/JNEUROSCI.0524-18.2018.

Ben-Yakov, A., Eshel, N., and Dudai, Y. (2013). Hippocampal immediate poststimulus activity in the encoding of consecutive naturalistic episodes. J. Exp. Psychol. Gen. 142, 1255–1263. https://doi.org/10.1037/a0033558.

Benjamini, Y., and Yekutieli, D. (2001). The control of the false discovery rate in multiple testing under dependency. Ann. Stat. https://doi.org/10.1214/aos/1013699998.

Bird, C.M. (2020). How do we remember events? Curr. Opin. Behav. Sci. 32, 120–125. https://doi.org/10.1016/j.cobeha.2020.01.020.

Braga, R.M., and Buckner, R.L. (2017). Parallel Interdigitated Distributed Networks within the Individual Estimated by Intrinsic Functional Connectivity. Neuron 95, 457–471.e5. https://doi.org/10.1016/j.neuron.2017.06.038.

Braga, R.M., Van Dijk, K.R.A., Polimeni, J.R., Eldaief, M.C., and Buckner, R.L. (2019). Parallel distributed networks resolved at high resolution reveal close juxtaposition of distinct regions. J. Neurophysiol. 121, 1513–1534. https://doi.org/10.1152/jn.00808.2018.

Brunec, I.K., Bellana, B., Ozubko, J.D., Man, V., Robin, J., Liu, Z.-X., Grady, C., Rosenbaum, R.S., Winocur, G., Barense, M.D., et al. (2017). Differential spatiotemporal representations along the hippocampal long axis in humans. BioRxiv.

Chen, J., Leong, Y.C., Honey, C.J., Yong, C.H., Norman, K.A., and Hasson, U. (2017). Shared memories reveal shared structure in neural activity across individuals. Nat. Neurosci. 20, 115–125. https://doi.org/10.1038/nn.4450.

Cohen, N.J., Poldrack, R.A., and Eichenbaum, H. (1997). Memory for Items and Memory for Relations in the Procedural/Declarative Memory Framework. Memory 5, 131–178. https://doi.org/10.1080/741941149.

Cohn-Sheehy, B.I., Delarazan, A.I., Reagh, Z.M., Crivelli-Decker, J.E., Kim, K., Barnett, A.J., Zacks, J.M., and Ranganath, C. (2021). The hippocampus constructs narrative memories across distant events Highlights. Curr. Biol. 1–11. https://doi.org/https://doi.org/10.1016/j.cub.2021.09.013.

Cooper, R.A., Kurkela, K.A., Davis, S.W., and Ritchey, M. (2021). Mapping the organization and dynamics of the posterior medial network during movie watching. Neuroimage 236, 118075. https://doi.org/10.1016/j.neuroimage.2021.118075.

Cox, R.W., and Hyde, J.S. (1997). Software tools for analysis and visualization of fMRI data. NMR Biomed. 10, 171–178. https://doi.org/10.1002/(sici)1099-1492(199706/08)10:4/5<171::aid-nbm453>3.0.co;2-l.

De Curtis, M., and Paré, D. (2004). The rhinal cortices: A wall of inhibition between the neocortex and the hippocampus. Prog. Neurobiol. 74, 101–110. https://doi.org/10.1016/j.pneurobio.2004.08.005.

Dale, A.M., Fischl, B., and Sereno, M.I. (1999). Cortical surface-based analysis. I. Segmentation and surface reconstruction. Neuroimage 9, 179–194. https://doi.org/10.1006/nimg.1998.0395.

DiNicola, L.M., Braga, R.M., and Buckner, R.L. (2020). Parallel distributed networks dissociate episodic and social functions within the individual. J. Neurophysiol. 123, 1144–1179. https://doi.org/10.1152/jn.00529.2019.

Ebbinghaus, H. (1885). Memory: A contribution to experimental psychology (translated by HA ruger & CE bussenues, 1913). New York Teach. Coll. Columbia Univ.

Finn, E.S. (2021). Is it time to put rest to rest? Trends Cogn. Sci. 1–12. https://doi.org/10.1016/j.tics.2021.09.005.

Fisher, J.S., and Radvansky, G.A. (2018). Patterns of forgetting. J. Mem. Lang. 102, 130–141. https://doi.org/10.1016/j.jml.2018.05.008.

Fonov, V., Evans, A., McKinstry, R., Almli, C., and Collins, D. (2009). Unbiased nonlinear average age-appropriate brain templates from birth to adulthood. Neuroimage 47, S102. https://doi.org/10.1016/S1053-8119(09)70884-5.

Frankland, P.W., and Bontempi, B. (2005). The organization of recent and remote memories. Nat. Rev. Neurosci. 6, 119–130. https://doi.org/10.1038/nrn1607.

Franklin, N.T., Norman, K.A., Ranganath, C., Zacks, J.M., and Gershman, S.J. (2020). Structured Event Memory: A neuro-symbolic model of event cognition. Psychol. Rev. 127, 327– 361. https://doi.org/10.1037/rev0000177.

Geerligs, L., Van Gerven, M., Campbell, K.L., and Güçlü, U. (2021). A nested cortical hierarchy of neural states underlies event segmentation in the human brain. BioRxiv 2021.02.05.429165. .

Ghosh, V.E., and Gilboa, A. (2014). What is a memory schema? A historical perspective on current neuroscience literature. Neuropsychologia 53, 104–114. https://doi.org/10.1016/j.neuropsychologia.2013.11.010.

Glasser, M.F., Coalson, T.S., Robinson, E.C., Hacker, C.D., Harwell, J., Yacoub, E., Ugurbil, K., Andersson, J., Beckmann, C.F., Jenkinson, M., et al. (2016). A multi-modal parcellation of human cerebral cortex. Nature 108, 125–138. https://doi.org/10.1038/nature18933.

Gordon, E.M., Laumann, T.O., Gilmore, A.W., Newbold, D.J., Greene, D.J., Berg, J.J., Ortega, M., Hoyt-Drazen, C., Gratton, C., Sun, H., et al. (2017a). Precision Functional Mapping of Individual Human Brains. Neuron 95, 791–807.e7. https://doi.org/10.1016/j.neuron.2017.07.011.

Gordon, E.M., Laumann, T.O., Adeyemo, B., and Petersen, S.E. (2017b). Individual Variability of the System-Level Organization of the Human Brain. Cereb. Cortex 27, 386–399. https://doi.org/10.1093/cercor/bhv239.

Gordon, E.M., Laumann, T.O., Marek, S., Raut, R. V., Gratton, C., Newbold, D.J., Greene, D.J., Coalson, R.S., Snyder, A.Z., Schlaggar, B.L., et al. (2020). Default-mode network streams for coupling to language and control systems. Proc. Natl. Acad. Sci. U. S. A. 117, 17308–17319. https://doi.org/10.1073/pnas.2005238117.

Gorgolewski, K., Burns, C.D., Madison, C., Clark, D., Halchenko, Y.O., Waskom, M.L., and Ghosh, S.S. (2011). Nipype: A Flexible, Lightweight and Extensible Neuroimaging Data Processing Framework in Python. Front. Neuroinform. 5. https://doi.org/10.3389/fninf.2011.00013.

Greve, D.N., and Fischl, B. (2009). Accurate and robust brain image alignment using boundary based registration. Neuroimage 48, 63–72. https://doi.org/10.1016/j.neuroimage.2009.06.060.

Griffiths, B.J., and Fuentemilla, L. (2020). Event conjunction: How the hippocampus integrates episodic memories across event boundaries. Hippocampus 30, 162–171. https://doi.org/10.1002/hipo.23161.

Hasson, U., Chen, J., and Honey, C.J. (2015). Hierarchical process memory: Memory as an integral component of information processing. Trends Cogn. Sci. 19, 304–313. https://doi.org/10.1016/j.tics.2015.04.006.

Heusser, A.C., Poeppel, D., Ezzyat, Y., and Davachi, L. (2016). Episodic sequence memory is supported by a theta-gamma phase code. Nat. Neurosci. 19, 1374–1380. https://doi.org/10.1038/nn.4374.

Howard, M.W., Fotedar, M.S., Datey, A. V., and Hasselmo, M.E. (2005). The Temporal Context Model in Spatial Navigation and Relational Learning: Toward a Common Explanation of Medial Temporal Lobe Function Across Domains. Psychol. Rev. 112, 75–116. https://doi.org/10.1037/0033-295X.112.1.75.

Hutchison, R.M., Womelsdorf, T., Allen, E.A., Bandettini, P.A., Calhoun, V.D., Corbetta, M., Della Penna, S., Duyn, J.H., Glover, G.H., Gonzalez-Castillo, J., et al. (2013). Dynamic functional connectivity: Promise, issues, and interpretations. Neuroimage 80, 360–378. https://doi.org/10.1016/j.neuroimage.2013.05.079.

Jenkinson, M., Bannister, P., Brady, M., and Smith, S. (2002). Improved optimization for the robust and accurate linear registration and motion correction of brain images. Neuroimage 17, 825–841. https://doi.org/10.1016/S1053-8119(02)91132-8.

Kahn, I., Andrews-Hanna, J.R., Vincent, J.L., Snyder, A.Z., and Buckner, R.L. (2008). Distinct cortical anatomy linked to subregions of the medial temporal lobe revealed by intrinsic functional connectivity. J. Neurophysiol. 100, 129–139. https://doi.org/10.1152/jn.00077.2008.

van Kesteren, M.T.R., Ruiter, D.J., Fernández, G., and Henson, R.N. (2012). How schema and novelty augment memory formation. Trends Neurosci. 35, 211–219. https://doi.org/10.1016/j.tins.2012.02.001.

Keven, N., Kurczek, J., Rosenbaum, R.S., and Craver, C.F. (2018). Narrative construction is intact in episodic amnesia. Neuropsychologia 110, 104–112. https://doi.org/10.1016/j.neuropsychologia.2017.07.028.

Klein, A., Ghosh, S.S., Bao, F.S., Giard, J., Häme, Y., Stavsky, E., Lee, N., Rossa, B., Reuter, M., Chaibub Neto, E., et al. (2017). Mindboggling morphometry of human brains. PLOS Comput. Biol. 13, e1005350. https://doi.org/10.1371/journal.pcbi.1005350.

Koen, J.D., Borders, A.A., Petzold, M.T., and Yonelinas, A.P. (2017). Visual short-term memory for high resolution associations is impaired in patients with medial temporal lobe damage. Hippocampus 27, 184–193. https://doi.org/10.1002/hipo.22682.

Kolarik, B.S., Baer, T., Shahlaie, K., Yonelinas, A.P., and Ekstrom, A.D. (2018). Close but no cigar: Spatial precision deficits following medial temporal lobe lesions provide novel insight into theoretical models of navigation and memory. Hippocampus 28, 31–41. https://doi.org/10.1002/hipo.22801.

Lanczos, C. (1964). Evaluation of noisy data. J. Soc. Ind. Appl. Math. Ser. B Numer. Anal. 1, 76–85. .

Lee, H., Bellana, B., and Chen, J. (2020). What can narratives tell us about the neural bases of human memory? Curr. Opin. Behav. Sci. 32, 111–119. https://doi.org/10.1016/j.cobeha.2020.02.007.

Levine, B., Svoboda, E., Hay, J.F., Winocur, G., and Moscovitch, M. (2002). Aging and autobiographical memory: dissociating episodic from semantic retrieval. Psychol. Aging 17, 677.

Lin, X., Sur, I., Nastase, S.A., Divakaran, A., Hasson, U., and Amer, M.R. (2019). Data-Efficient Mutual Information Neural Estimator. 1–16.

Lu, Q., Hasson, U., and Norman, K.A. (2022). A neural network model of when to retrieve and encode episodic memories. Elife 11, 1–46. https://doi.org/10.7554/elife.74445.

McClelland, J.L., McNaughton, B.L., and O’Reilly, R.C. (1995). Why there are complementary learning systems in the hippocampus and neocortex: Insights from the successes and failures of connectionist models of learning and memory. Psychol. Rev. 102, 419–457. https://doi.org/10.1037/0033-295X.102.3.419.

Michelmann, S., Price, A.R., Aubrey, B., Strauss, C.K., Doyle, W.K., Friedman, D., Dugan, P.C., Devinsky, O., Devore, S., Flinker, A., et al. (2021). Moment-by-moment tracking of naturalistic learning and its underlying hippocampo-cortical interactions. Nat. Commun. 12, 1–15. https://doi.org/10.1038/s41467-021-25376-y.

Milivojevic, B., Varadinov, M., Grabovetsky, A.V., Collin, S.H.P., and Doeller, C.F. (2016). Coding of event nodes and narrative context in the hippocampus. J. Neurosci. 36, 12412–12424. https://doi.org/10.1523/JNEUROSCI.2889-15.2016.

Mukamel, R., Gelbard, H., Arieli, A., Hasson, U., Fried, I., and Malach, R. (2005). Coupling Between Neuronal Firing, Field Potentials, and fMRI in Human Auditory Cortex. Science (80-.). 309, 951–954. https://doi.org/10.1126/science.1110913.

Nadel, L., and Moscovitch, M. (1997). Memory consolidation, retrograde amnesia and the hippocampal complex. Curr. Opin. Neurobiol. 7, 217–227. https://doi.org/10.1016/S0959-4388(97)80010-4.

O’Reilly, R.C., Ranganath, C., and Russin, J.L. (2021). The Structure of Systematicity in the Brain.

O’Reilly, R.C., Ranganath, C., and Russin, J.L. (2022). The Structure of Systematicity in the Brain. Curr. Dir. Psychol. Sci. 096372142110492. https://doi.org/10.1177/09637214211049233.

Oedekoven, C.S.H., Keidel, J.L., Berens, S.C., and Bird, C.M. (2017). Reinstatement of memory representations for lifelike events over the course of a week. Sci. Rep. 7, 1–12. https://doi.org/10.1038/s41598-017-13938-4.

Oedekoven, C.S.H., Keidel, J.L., Anderson, S., Nisbet, A., and Bird, C.M. (2019). Effects of amnesia on processing in the hippocampus and default mode network during a naturalistic memory task: A case study. Neuropsychologia 132, 107104. https://doi.org/10.1016/j.neuropsychologia.2019.05.022.

Peirce, J., Gray, J.R., Simpson, S., MacAskill, M., Höchenberger, R., Sogo, H., Kastman, E., and Lindeløv, J.K. (2019). PsychoPy2: Experiments in behavior made easy. Behav. Res. Methods 51, 195–203. https://doi.org/10.3758/s13428-018-01193-y.

Poppenk, J., and Moscovitch, M. (2011). A hippocampal marker of recollection memory ability among healthy young adults: contributions of posterior and anterior segments. Neuron 72, 931–937. https://doi.org/10.1016/j.neuron.2011.10.014.

Poppenk, J., Evensmoen, H.R., Moscovitch, M., and Nadel, L. (2013). Long-axis specialization of the human hippocampus. Trends Cogn. Sci. 17, 230–240. https://doi.org/10.1016/j.tics.2013.03.005.

Power, J.D., Mitra, A., Laumann, T.O., Snyder, A.Z., Schlaggar, B.L., and Petersen, S.E. (2014). Methods to detect, characterize, and remove motion artifact in resting state fMRI. Neuroimage 84, 320–341. https://doi.org/10.1016/j.neuroimage.2013.08.048.

Preston, A.R., and Eichenbaum, H. (2013). Interplay of hippocampus and prefrontal cortex in memory. Curr. Biol. 23, R764–R773. https://doi.org/10.1016/j.cub.2013.05.041.

Radvansky, G.A., and Zacks, J.M. (2014). Event cognition (Oxford University Press).

Ranganath, C., and Ritchey, M. (2012). Two cortical systems for memory-guided behaviour. Nat. Rev. Neurosci. 13, 713–726. https://doi.org/10.1038/nrn3338.

Reagh, Z.M., and Ranganath, C. (2018). What does the functional organization of cortico-hippocampal networks tell us about the functional organization of memory? Neurosci. Lett. 680, 69–76. https://doi.org/10.1016/j.neulet.2018.04.050.

Reagh, Z.M., and Ranganath, C. (2021). A cortico-hippocampal scaffold for representing and recalling lifelike events. BioRxiv 2021.04.16.439894. https://doi.org/10.1101/2021.04.16.439894.

Reagh, Z.M., Delarazan, A.I., Garber, A., and Ranganath, C. (2020). Aging alters neural activity at event boundaries in the hippocampus and Posterior Medial network. Nat. Commun. 11, 1–12. https://doi.org/10.1038/s41467-020-17713-4.

Reuter, M., Rosas, H.D., and Fischl, B. (2010). Highly accurate inverse consistent registration: A robust approach. Neuroimage 53, 1181–1196. https://doi.org/10.1016/j.neuroimage.2010.07.020.

Ritchey, M., and Cooper, R.A. (2020). Deconstructing the Posterior Medial Episodic Network. Trends Cogn. Sci. 1–15. https://doi.org/10.1016/j.tics.2020.03.006.

Robin, J., and Moscovitch, M. (2017). Details, gist and schema: hippocampal–neocortical interactions underlying recent and remote episodic and spatial memory. Curr. Opin. Behav. Sci. 17, 114–123. https://doi.org/10.1016/j.cobeha.2017.07.016.

Rugg, M.D., and Vilberg, K.L. (2013). Brain networks underlying episodic memory retrieval. Curr. Opin. Neurobiol. 23, 255–260. https://doi.org/10.1016/j.conb.2012.11.005.

Rumelhart, D.E., and Ortony, A. (1977). The Representation of Knowledge in Memory. In Schooling and the Acquisition of Knowledge, (Routledge), pp. 99–135.

Sekeres, M.J., Bonasia, K., St-Laurent, M., Pishdadian, S., Winocur, G., Grady, C., and Moscovitch, M. (2016). Recovering and preventing loss of detailed memory: Differential rates of forgetting for detail types in episodic memory. Learn. Mem. 23, 72–82. https://doi.org/10.1101/lm.039057.115.

Sekeres, M.J., Winocur, G., Moscovitch, M., Anderson, J.A.E., Pishdadian, S., Martin Wojtowicz, J., St-Laurent, M., McAndrews, M.P., and Grady, C.L. (2018). Changes in patterns of neural activity underlie a time-dependent transformation of memory in rats and humans. Hippocampus 28, 745–764. https://doi.org/10.1002/hipo.23009.

Silva, M., Baldassano, C., and Fuentemilla, L. (2019). Rapid Memory Reactivation at Movie Event Boundaries Promotes Episodic Encoding. J. Neurosci. 39, 8538–8548. https://doi.org/10.1523/JNEUROSCI.0360-19.2019.

Squire, L.R., and Zola-Morgan, S. (1991). The medial temporal lobe memory system. Science 253, 1380–1386. .

Strange, B.A., Witter, M.P., Lein, E.S., and Moser, E.I. (2014). Functional organization of the hippocampal longitudinal axis. Nat. Rev. Neurosci. 15, 655–669. https://doi.org/10.1038/nrn3785.

Teyler, T.J., and DiScenna, P. (1986). The Hippocampal Memory Indexing Theory. Behav. Neurosci. 100, 147–154. https://doi.org/10.1037/0735-7044.100.2.147.

Thakral, P.P., Madore, K.P., and Schacter, D.L. (2020). The core episodic simulation network dissociates as a function of subjective experience and objective content. Neuropsychologia 136, 107263. https://doi.org/10.1016/j.neuropsychologia.2019.107263.

Thorndyke, P.W. (1977). Cognitive structures in comprehension and memory of narrative discourse. Cogn. Psychol. 9, 77–110. https://doi.org/10.1016/0010-0285(77)90005-6.

Tustison, N.J., Avants, B.B., Cook, P.A., Yuanjie Zheng, Egan, A., Yushkevich, P.A., and Gee, J.C. (2010). N4ITK: Improved N3 Bias Correction. IEEE Trans. Med. Imaging 29, 1310–1320. https://doi.org/10.1109/TMI.2010.2046908.

Whitfield-Gabrieli, S., and Nieto-Castanon, A. (2012). Conn: a functional connectivity toolbox for correlated and anticorrelated brain networks. Brain Connect. 2, 125–141. https://doi.org/10.1089/brain.2012.0073.

Zadbood, A., Chen, J., Leong, Y.C., Norman, K.A., and Hasson, U. (2017). How We Transmit Memories to Other Brains: Constructing Shared Neural Representations Via Communication. Cereb. Cortex 27, 4988–5000. https://doi.org/10.1093/cercor/bhx202.

Zhang, Y., Brady, M., and Smith, S. (2001). Segmentation of brain MR images through a hidden Markov random field model and the expectation-maximization algorithm. IEEE Trans. Med. Imaging 20, 45–57. https://doi.org/10.1109/42.906424.

Zheng, J., Schjetnan, A.G.P., Yebra, M., Gomes, B.A., Mosher, C.P., Kalia, S.K., Valiante, T.A., Mamelak, A.N., and Kreiman, G. (2022). Neurons detect cognitive boundaries to structure episodic memories in humans. 25. https://doi.org/10.1038/s41593-022-01020-w.

Zuo, X., Honey, C.J., Barense, M.D., Crombie, D., Norman, K.A., Hasson, U., and Chen, J. (2020). Temporal integration of narrative information in a hippocampal amnesic patient. Neuroimage 213, 116658. https://doi.org/10.1016/j.neuroimage.2020.116658.

