## Supplemental Methods and Results for "Dynamic hippocampal-cortical interactions during event boundaries support retention of complex narrative events"

#### Recall statistical analysis

A mixed linear model was used to determine the effect of delay on the number of events subsequently recalled, and the number of details recalled. An event was considered recalled if participants mentioned any detail that was specific to a given event. Movie and counter-balance group were entered as covariates of no interest. The model predicting number of details recalled from delay also included detail type (central vs. peripheral) as an effect of interest. The Shapiro–Wilk test was used to evaluate normality on the distribution of details.

### Supplemental Results

#### Recall statistical analysis

Participants retrieved a fewer number of events after a 2-day delay compared to the immediate recall condition ($t(44) = 4.91$, $p < .001$). The distribution of the number of details recalled by participants was skewed, with some subjects recalling many more details above average than the group. A Shapiro-Wilk test revealed that the distribution was not normal, $W = .72$, $p < .001$. The distribution was therefore log-transformed using the natural log, resulting in a normal distribution, $W = 0.97$, $p = .058$. Using the log-transformed detail scoring, we observed that participants reported fewer details after a 2-day delay compared to the immediate recall condition ($t(88) = 3.2$, $p = .001$). Participants also tended to report more peripheral details than central details ($t(88) = 8.7$, $p < .001$), but there was no interaction between detail type and delay in terms of details recalled ($t(88) = .36$, $p = .7$. In all models, there was no effect of movie, (t-values < 1, p-values > 0.29) or counter-balance group, (F-values < 1.76, p-values > .17). Since there were no interactions with detail type, future analysis pooled details into total details.

Supplemental Figures

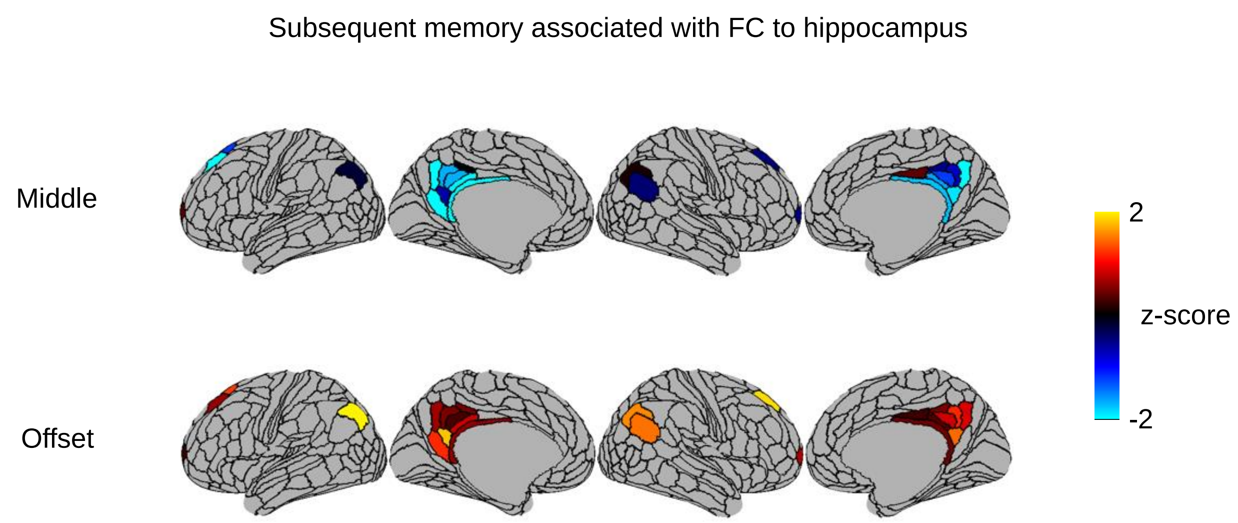


**Supplemental Figure 1**

Relationship between subsequent recall success and hippocampal FC with ROIs in the PMN plotted on an inflated brain surface. Warm colors indicate that functional connectivity between the ROI and hippocampus is associated with better subsequent memory for events and cool colors indicate that functional connectivity with the hippocampus is associated with worse subsequent memory.

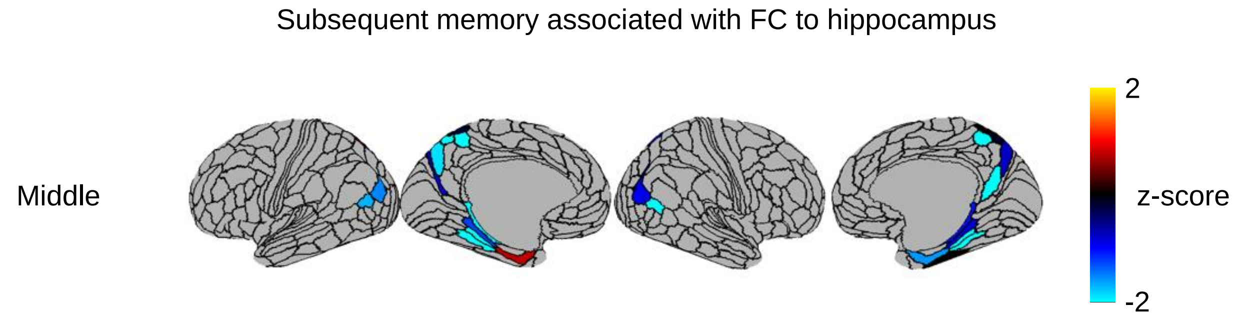


**Supplemental Figure 2**

Relationship between subsequent recall success and hippocampal FC with ROIs in the MTN at the immediate recall time point plotted on an inflated brain surface.

Details associated with FC to hippocampus

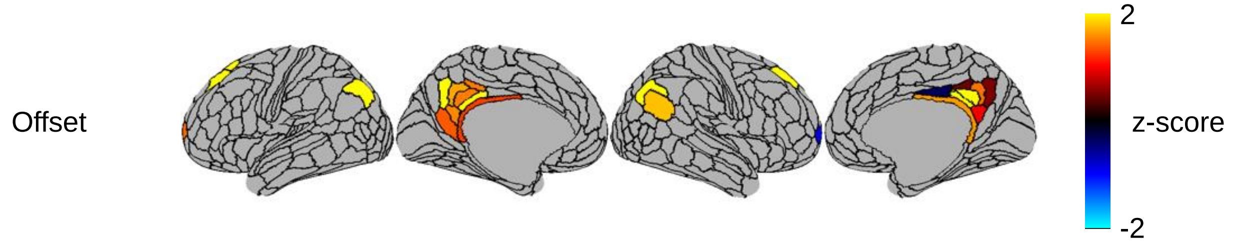


**<u>Supplemental Figure 3</u>**

Relationship between number of subsequently recalled and hippocampal FC with ROIs in the PMN at the delayed recall time point plotted on an inflated brain surface.
